# The Behaviour of F-statistics over Time

**DOI:** 10.1101/2022.08.25.505252

**Authors:** Song Li, Carsten Wiuf

**Affiliations:** Department of Mathematical Sciences, University of Copenhagen, Denmark

**Keywords:** Allele frequency, *F*-statistics, Wright–Fisher model, Linear evolutionary force, Finite population, Population genetics

## Abstract

We study the behaviour of the *F*_2_-statistic and *F*_st_-statistic, respectively, over time in a Wright-Fisher model with mutation and migration. We give precise conditions for when the *F*_2_-statistic is non-monotonic, that is, increases over time until a certain point and then starts decreasing. We show that even for small population sizes, the two statistics are well approximated by population size scaled expressions.

## 1 Introduction

The frequency of an allele in a population depends on various evolutionary forces. The Wright–Fisher model (Fisher, 1930; Wright, 1931) in its simplest form describes the evolution of the allele frequency at a single diploid site in a finite population, without overlapping generations, in the absence of any other evolutionary forces such as mutation and selection. For large populations sizes, using that the Wright-Fisher model can be regarded as a discrete time Markov process, continuous time diffusion approximations can be derived by scaling time and parameters in the population size (Crow and Kimura, 1970; Ewens, 2004; Karlin and Taylor, 1980; Neuhauser, 2001). The diffusion process is essentially determined by its first two moments, which greatly simplifies the exploration of the behavior of the allele frequency over time in large populations. Recently, other approximations have been proposed to study the distribution of the allele frequency over finitely many generations, based on the Wright-Fisher model, e.g., Balding and Nichols (1995); Nicholson et al. (2002); Foll and Gaggiotti. (2008); Coop et al. (2010); Gautier (2015); Haasl and Payseur (2016).

The main purpose of this paper is to explore the behavior of the two *F* -statistics, *F*_2_ and *F*_*st*_, that reflect frequency changes and population differentiation. These statistics depend on the first two moments of the allele frequency distribution. For multiple populations, *F*_2_ and *F*_*st*_ are related (Reich et al., 2009; Peter, 2016). The *F*_*st*_, or the fixation index, is a measurement of population differences in alelle frequency and can be defined in two ways (Holsinger and Weir, 2009; Durrett, 2008). The *F*_*st*_-statistic provides important insights into frequency processes within and between populations. Pure drift considered in the Wright-Fisher process is the simplest evolutionary force. However, evolution is complex and random, involving other factors such as mutation, migration, and natural selection (Cavalli-Sforza and Edwards, 1967). In general, evolutionary forces are divided into linear and nonlinear forms (Crow and Kimura, 1970). Linear forces are typically mutation and migration, which we also consider here. In this paper, we adoopt the definition of *F*_2_ proposed by Reich et al. (2009), and the definition of *F*_*st*_ proposed by (Wright, 1951). The *F*_2_ is defined as the square of the difference in allele frequency between two populations and has a range of mathematical properties (such as additivity) used in admixture inference (Patterson et al., 2012; Peter, 2016; Soraggi and Wiuf, 2019). Here, we study how *F*_2_ and *F*_*st*_ vary over time. In particular, we show that migration might give rise to non-monotome behavior and analyse when this happens. We give precise conditions for when an inflection point occurs, that is, a time point after which the statisitc starts decreasing. Under pure drift, both statistics increase over time.

The paper is structured as follows. We describe the Wright-Fisher model in Section 2.1, allowing for mutation and migration. In Section 2.2, we give the definition of the *F*_2_-statistic and the *F*_*st*_-statistic, and find expressions for how they vary over time. We end with a discussion in Section 3. Proofs and mathematical details are collected in the Appendix.

## 2 Methods and Results

We consider a population of haploid individuals over generations. The frequency of the reference allele is set to *X*_*t*_ ∈ [0, 1] in generation *t* ≥ 0, and we are interested in the evolution of *X*_*t*_ over time. On this basis, we impose certain constraints on *X*_*t*_, *t* ≥ 0, to establish a model for its change.

Specifically, we consider a Wright-Fisher-like model with population size *N*_*t*_ in generation *t* ≥ 0. The random number of individuals carrying the reference allele is denoted *Z*_*t*_, hence *X*_*t*_ = *Z*_*t*_*/N*_*t*_. In line with the Wirght-Fisher model, we assume the allele frequency of the next generation is determined by the previous generation and potentially external factors.

The external influence acting on the allele can be defined as a function *g* : [0, 1] → [0, 1], and the model can be set up using a binomial distribution,

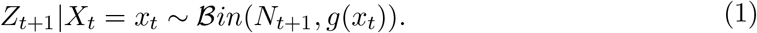

Here, ℬ*in*(*n, p*) is the binomial distribution with sample size *n* and probability *p*. The mathematical form of the effect *g* will be discussed below. The Wright-Fisher model can be extended to diploids by changing *N* to 2*N*, or to non-constant population sizes by taking floating samples from each generation.

### 2.1 Linear evolutionary pressure model

We are mainly concerned with the influence of pure drift, mutation and migration on the evolution of the allele frequency (Cavalli-Sforza and Edwards, 1967; Cavalli-Sforza, 1973). The common characteristic is a linear constraint on how the frequency change (Siren, 2012). Specifically, we address the following cases.

In the case of mutation, *a*_1_ is assumed to be the probability of mutation from the allele ‘A’ to the allele ‘a’, and *b*_1_ is the probability from ‘a’ to ‘A’. Therefore, if the frequency of ‘A’ carried by the parent is *X*_*t*_, then we define

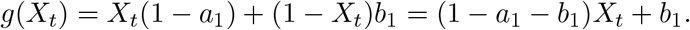

In the simplest case of migration (Tataru et al., 2015, 2016), individuals have the freedom to migrate in and out of the population. Assume that the probability of migration between populations is *m* and an infinitely large population with a constant allele frequency *X*^*^ ∈ [0, 1]. Then, we get the calculation

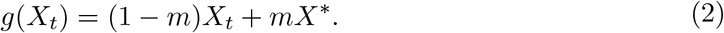

We incorporate both the above mutation and migration into the model,

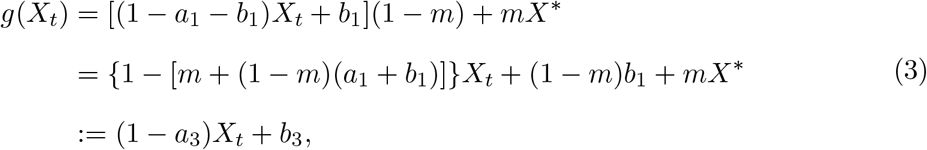

where, *a*_3_ = *m* + (1 − *m*)(*a*_1_ + *b*_1_), *b*_3_ = (1 − *m*)*b*_1_ + *mX*^*^. Obviously, by taking *X*_*t*_ to be 1, then 0 ≤ 1 − *a*_3_ + *b*_3_ ≤ 1, so we need 0 ≤ *b*_3_ ≤ *a*_3_ ≤ 1 as a constrain.

We describe the above results directly in terms of the function *g*,

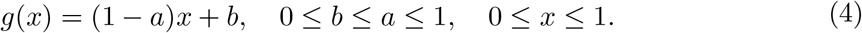

In particular, when *a* = *b* = 0, the above model degenerates into the familiar pure drift case.

### 2.2 Definitions of F-statistics

In this section, we study the *F*_2_ (Reich et al., 2009) and the *F*_st_ (Wright, 1951). The *F*_2_ can be used to measure the difference in allele frequencies in a single population at different time points, and *F*_*st*_ describes this difference in multiple populations. The following parts are our results.

#### 2.2.1 The F_2_

In a single population, we consider two time points 0 and *t*, and express their allele frequencies as *X*_0_ and *X*_*t*_, respectively. Then the *F*_2_ is defined as,

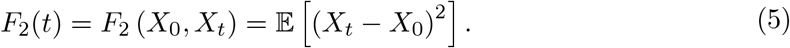

Since we are interested in the variation of the allele frequency from its starting point, we assume throughout that *X*_0_ = *x*_0_, with 0 ≤ *x*_0_ ≤ 1, is fixed. Then *F*_2_(*t*) becomes

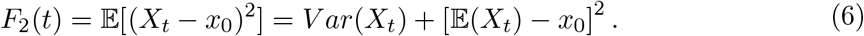

Under the linear evolutionary pressure model, let

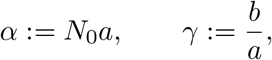

with the convention that 0*/*0 := 1, then 0 ≤ γ ≤ 1. The variance can be given recursively in terms of the expectation, similarly to Tataru et al. (2015, 2016) and Siren (2012),

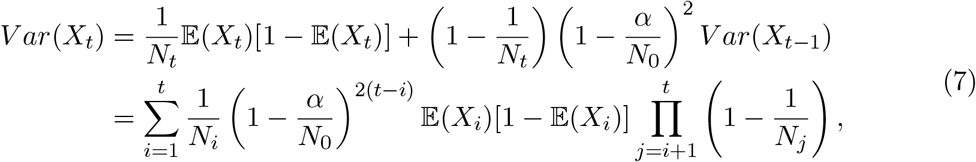

and

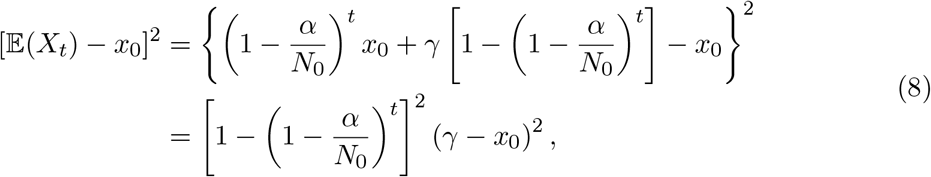

where

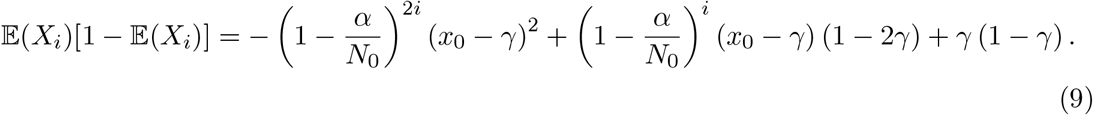

Equation (8) only depends on the population size through *N*_0_. In contrast, (7) depends on *N*_*t*_. Specifically, the effect of *N*_*t*_ is to locally slow down or speed up (compared to *N*_0_) the change in the variance: doubling the population size corresponds to slowing time by a factor of two, at least for large population sizes.

Furthermore, by scaling the generation (time) and the parameter *b* in units of *N*_0_, we introduce the notation,

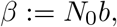

such that

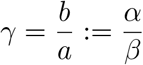

is independent of *N*_0_ and *t* = ⌊*N*_0_*u*⌋, where ⌊·⌋ is the rounding operation and *u* ∈ (0, ∞). We allow the population size to vary in a slow way, that is, we assume that there is a positive continuous function *h* : (0, ∞) → (0, ∞), such that

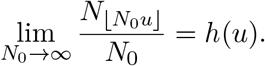

Such assumptions are widely used in population genetics, for example, to derive the diffusion limit of the Wright-Fisher model (Ewens, 2004).

Defining two limits 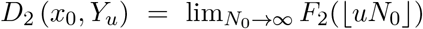 and 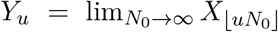 (Ewens, 2004), then we denote

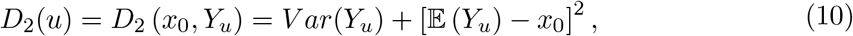

where, by defining 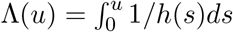 as the p opulation-size intensity f unction (Griffiths and Tavaré, 1994),

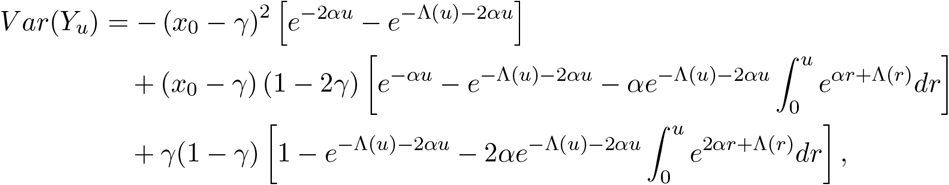

and

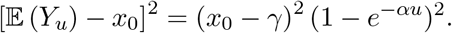

Through the above definitions, we obtain the explicit expression *F*_2_(*t*) in the general case and the limit expression *D*_2_(*u*) in the infinite size case. In the following part, we give some theoretical properties through the discussion of parameters.

#### 2.2.2 Properties for the F_2_

We first present results for a single population under pure drift in a population of any size. In pure drift case, *g*(*x*) = *x* and the expectation of frequency is 𝔼(*X*_*t*_) = 𝔼(*X*_*t*−1_) = *x*_0_.

Then

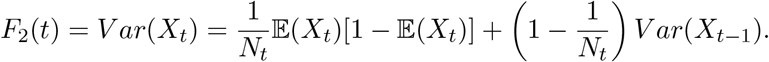

It follows that

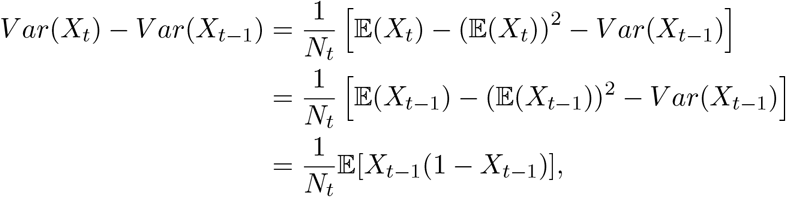

hence due to 0 ≤ *X*_*t*_ ≤ 1,

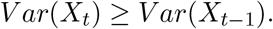

Proposition below gives the performance of *F*_2_(*t*) under pure drift.

##### Proposition.

*Under pure drift, the variance and F*_2_(*t*) *are gradually increasing over generations in a single population of any size (Barton and Turelli, 2004; Peter, 2016)*.

In the generous linear evolutionary pressure model, the change in *F*_2_(*t*) is not as straightforward as the case of pure drift. We assume *N*_*t*_ ≡ *N*_0_ = *N* to simplify the analysis.

Using (7), (8) and (9), we replace the expression for *F*_2_(*t*) in (6) by an expression without the recursion,

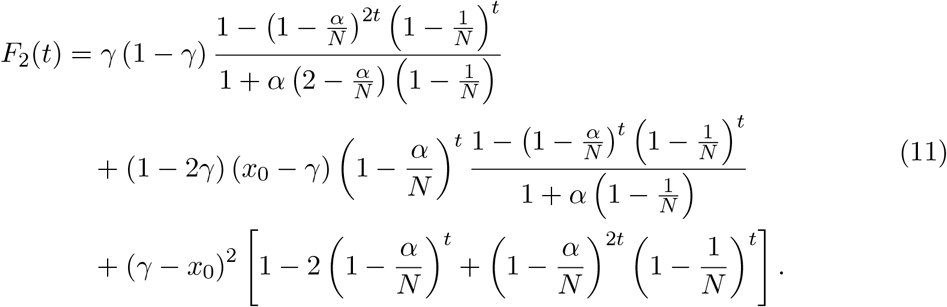

The expression above is consistent with the results in the literature (Siren, 2012; Tataru et al., 2015). It follows that the parameters (*x*_0_, γ, α, *N*) and (1 − *x*_0_, 1 − γ, α, *N*) result in the same *F*_2_(*t*) value. Thus, we might assume the parameter *x*_0_ lies in [0, 0.5].

Theorem 1 shows that *F*_2_(*t*) is not always increasing. Define Δ_1_ and Δ_2_ by

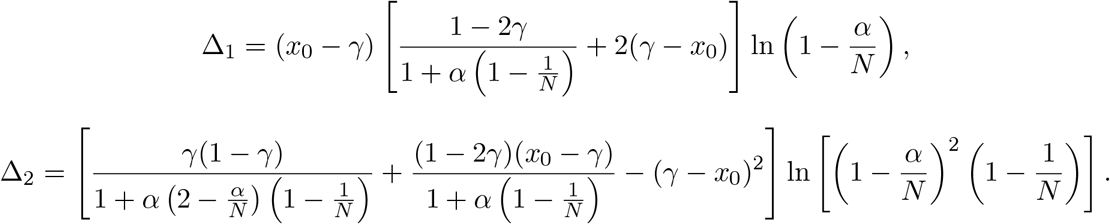

##### Theorem 1.

*Suppose N*_*t*_ ≡ *N*_0_ = *N, then F*_2_(*t*) *has an inflection point if and only if* Δ_1_ *<* 0, *and F*_2_(*t*) *is non-increasing for all* 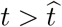, *where*

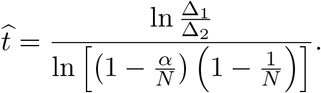

We replace the condition Δ_1_ *<* 0 by

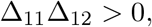

where,

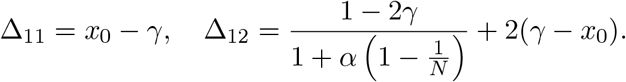

Thus, Δ_11_ and Δ_12_ must have the same sign for the condition to hold. According to this criterion, Figure 1 shows the region Δ_1_ *<* 0 for *x*_0_ = 0.2, 0.4 and *N* = 5, 10, 50, ∞. By allowing the population size to vary over time, clearly more complicated behaviors of *F*_2_(*t*) should be expected.

**Figure 1:**
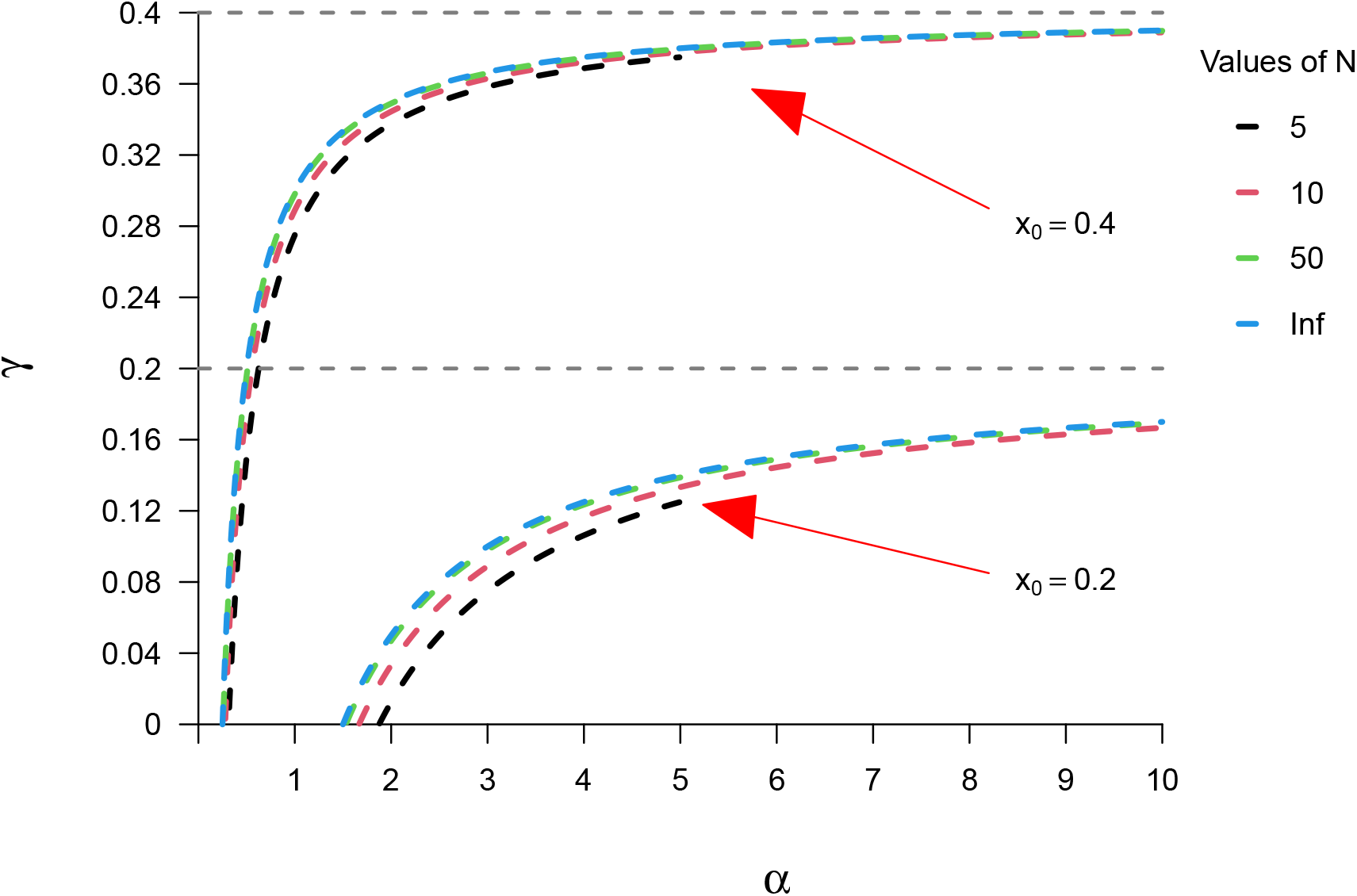
The region where Δ_1_ *<* 0. For *x*_0_ = 0.4, the region is divided by a gray horizontal line *γ* = 0.4, axis of coordinates and a series of color lines at the top left corner of the figure; for *x*_0_ = 0.2, the region is divided by a gray horizontal line *γ* = 0.2, axis of coordinates and a series of color lines at the bottom right corner of the figure.

Note that 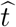 in Theorem 1 is defined as the inflection point of *F*_2_(*t*) from increase to decrease, whose visualization is analyzed in the following. Since the position of inflection point involves parameters *N, x*_0_, *α* and *γ*, in order to display the change of *F*_2_(*t*) intuitively, we choose fixed *N* = 100 and *x*_0_ = 0.2. By changing *α* and *γ*, we can give a heat map of inflection points, which shows the early and late appearance (see Figure 2).

**Figure 2:**
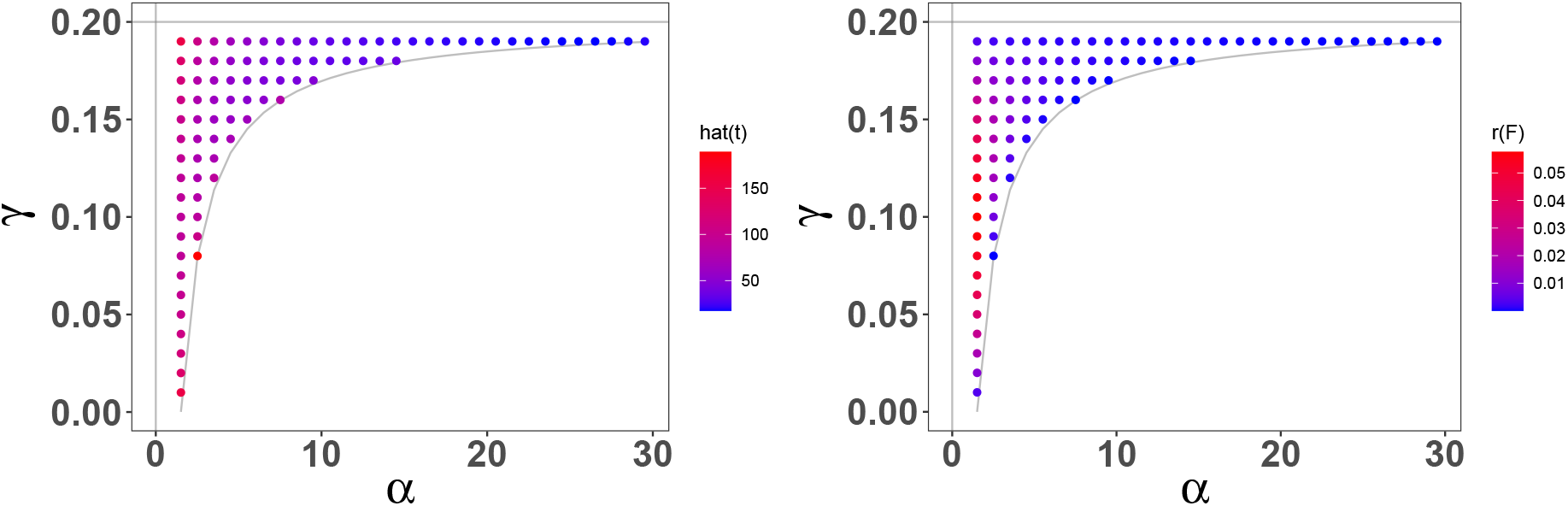
Shown is the heat map of 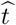 and *r*(*F*) under the condition of *N* = 100 and *x*

As shown in Figure 2, we choose the inflection point is near 100 and get the parameters *α* = 1.5158, *γ* = 0.12 that allow us to find the case where the inflection point should be (see Figure 3). As can be seen from Figure 3, 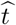 is marked as the moment when the variation (compared to *x*_0_) of population allele frequency is the greatest, and *t* at infinity represents a stationary state. The relative difference degree of *F*_2_(*t*) value between these two points (namely peak and plateau value) is also an indicator to understand the population genetic process. We give the following definition,

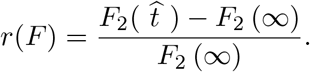

**Figure 3:**
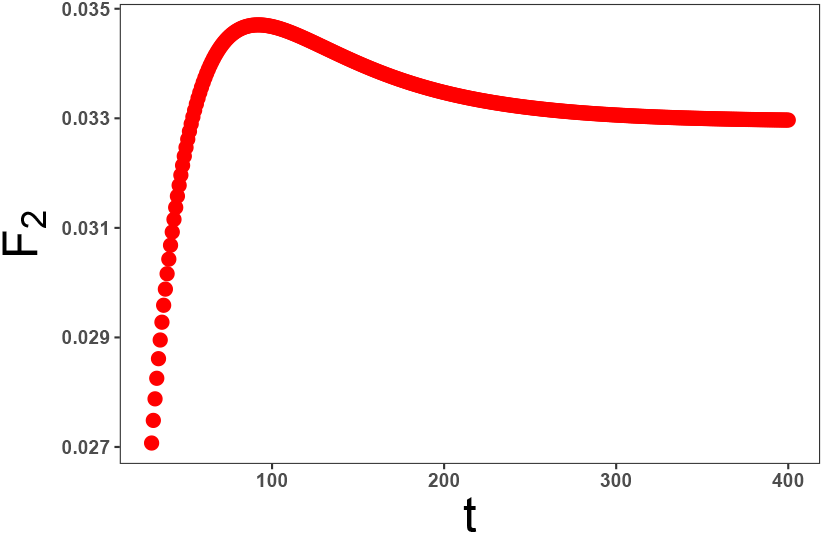
Shown is the population genetic evolution process with parameters *N* = 100, *x*_0_ = 0.2, *α* = 1.5158, and *γ* = 0.12. The *F*_2_ has obvious peak and plateau value.

Similarly, we show *r*(*F*) for different *α* and *γ* in the heat map (see Figure 2). In general, we find *r*(*F*) might be up to 0.1.

The case where *N* goes to infinity is defined as an infinite population model. In the infinite population case, we take some steps to get similar results. Under pure drift, we have

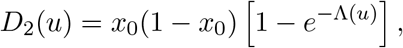

by setting *h* ≡ 1, *D*_2_(*u*) simplifies to

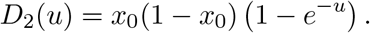

*D*_2_(*u*) in this case is increasing with *u*. In the generous linear evolutionary pressure model and by setting *h* ≡ 1, the expression (10) can be simplified to

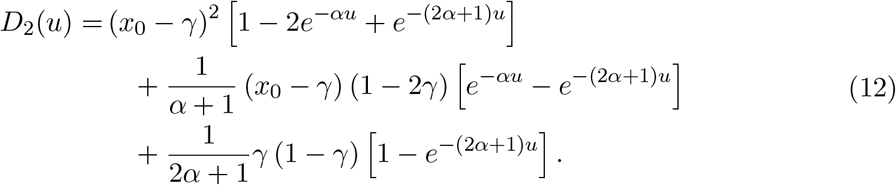

The parameters (*x*_0_, *γ, α*) and (1 − *x*_0_, 1 − *γ, α*) result in the same *D*_2_(*u*) value. Theorem 2 below gives a similar result to that of Theorem 1. Define Θ_1_ and Θ_2_ by

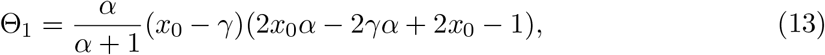

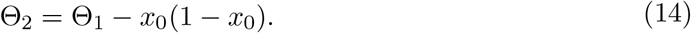

##### Theorem 2.

*Suppose h* ≡ 1, *then D*_2_(*u*) *has an inflection point if and only if* Θ_1_ *<* 0, *and if this is the case then D* (*u*) *is non-increasing for all u >* Û, *where*

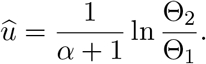

For consistency with the finite *N* case, we replace the condition Θ_1_ *<* 0 by

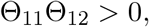

where

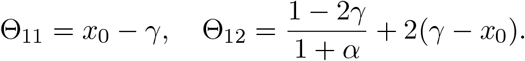

Thus, Θ_11_ and Θ_12_ must have the same sign for the condition to hold. According to this criterion, Figure 4 shows the region Θ_1_ *<* 0 for *x*_0_ = 0.2, 0.4, 0.5, 0.6 and 0.8.

**Figure 4:**
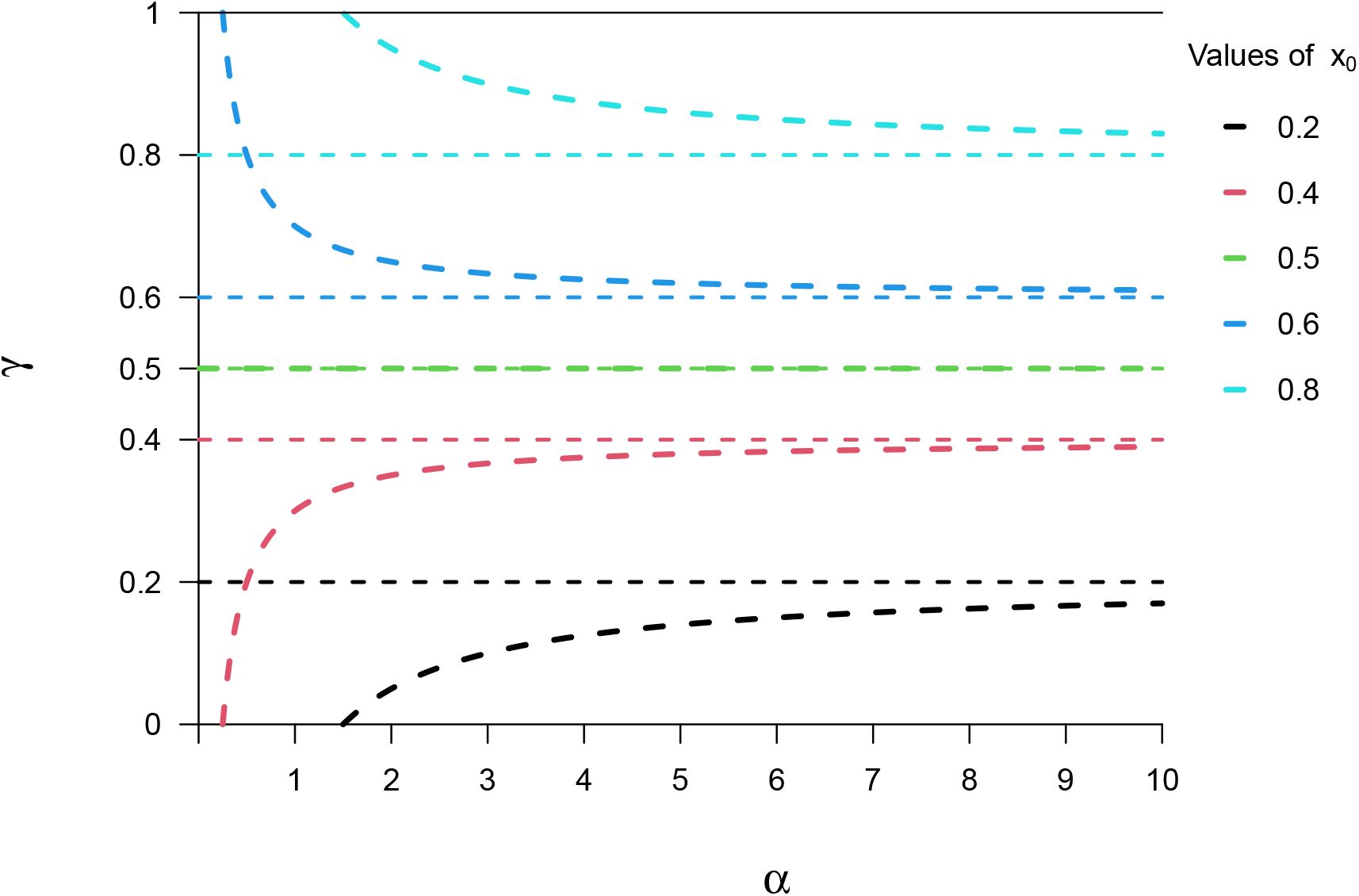
The region Θ_1_ *<* 0 for different values of *x*_0_ in the case *N* → ∞. For each given *x*_0_, the region is divided by a colored horizontal line *γ* = *x*_0_, a curve corresponding to the same color and axis of coordinates. For *x*_0_ = 0.5, the region is empty.

As shown in the Figure 4, if *x*_0_ = 0.5, then Θ_1_ ≥ 0, that means *D*_2_(*u*) keeps increasing for any *γ* and *α*. In contrast, the initial value *x*_0_ within a specified range can also determine that the inflection point must exist. The following corollary shows this and gives the specific location of the region.

##### Corollary.

*Let α* ≥ 0, *γ* ∈ [0, 1] *be given, such that γ* ≠ 0.5, *and let N be a natural number. Then F*_2_(*t*) *has an inflection point for any x*_0_ ∈ (*x*_0*L*_, *x*_0*R*_), *where*

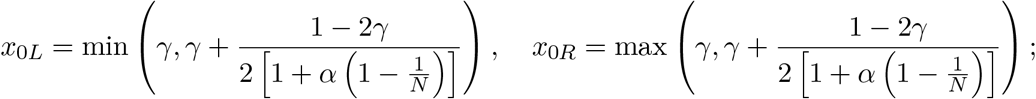

*Simialrly, D*_2_(*u*) *has an inflection point for any x*_0_ ∈ (*x*_0*L*_, *x*_0*R*_), *where*

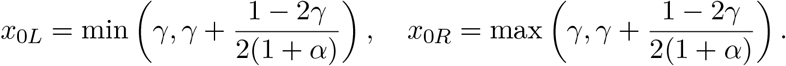

#### 2.2.3 The F_st_

For two or more populations, we study *F*_st_ which was first proposed by Sewall Wright (Wright, 1951) to measure the differences of allele frequencies among populations. Note that there are some different definitions of *F*_st_ (Nei, 1986; Holsinger and Weir, 2009; Durrett, 2008). For two different populations *P*_1_ and *P*_2_, we use the following definition of *F*_st_ in terms of probability,

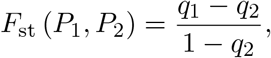

where *q*_1_ and *q*_2_ represent the probability that two given reference alleles are the same from within and between populations, respectively. Suppose that the reference allele has frequency 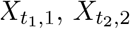 in population *P*_1_ with population size 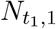 at time *t*_1_, and *P*_2_ with population size 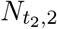 at time *t*_2_, respectively. Then, we define *q*_1_ and *q*_2_ as follows (Reich et al., 2009)

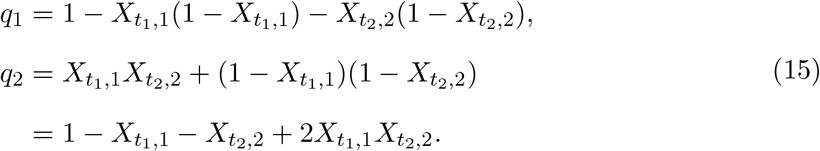

Using equation (15), we have

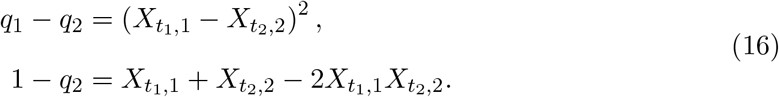

As *q*_1_, *q*_2_ are stochastic variables, we adopt the following definition, replacing *P*_1_, *P*_2_, with *t*_1_, *t*_2_, respectively,

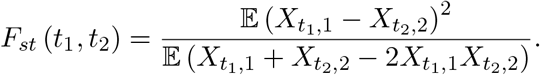

We regard *F*_st_ as a function that changes over time. For two different populations, if we only consider the linear evolutionary pressure model, the two branching populations from a common ancestral population are independent of each other in the subsequent evolutionary process (Hansen and Martins, 1996), i.e., 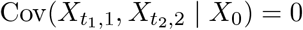. If *X*_0_ is fixed, indirectly, we have 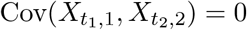.

In the following, we set 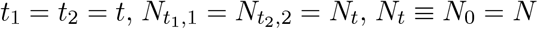, *N*_*t*_ ≡ *N*_0_ = *N* (indicating *h* ≡ 1), *F*_st_(*t*) = *F*_st_(*t, t*) and using the previous notation, we denote *F*_st_(*t*) as *D*_*st*_(*u*) when *N* is large. To facilitate the application of the above results, according to the definitions (6) and (10), we split *F*_*st*_(*t*) and *D*_*st*_(*t*) as follows,

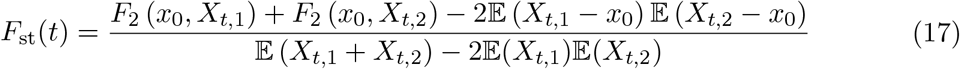

and

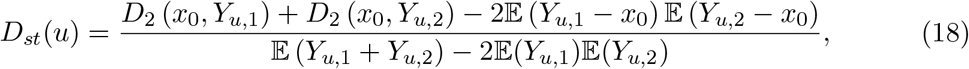

where *Y*_*u,i*_ = lim_*N*→∞_ *X*_⌊*uN* ⌋,*i*_, *i* = 1, 2.

If pure drift only is considered in the evolution of two populations, then expression (17) and (18) degenerate to

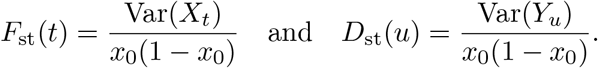

Based on the results for *F*_2_(*t*) (and *D*_2_(*u*)), then *F*_st_(*t*) (and *D*_st_(*u*)) is gradually increasing over generations for two different populations of any size under the pure drift. In the previous section, we elaborated on the properties of *F*_2_(*t*) and *D*_2_(*u*), which are consistent, and in the following, we only focus on the infinite population size case of *D*_st_(*u*) and the situation in which an inflection point occurs.

If we consider pure drift as the only evolutionary force factor for *P*_1_ and the general linear evolutionary pressure model for *P*_2_, then (18) degenerates to

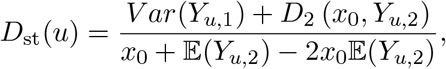

where,

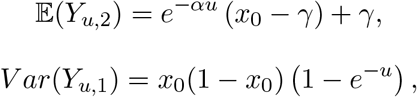

and using (13), (14), the expression (12) can be simplified to

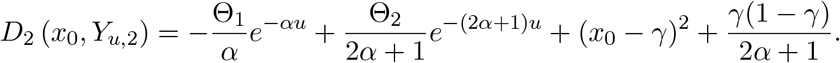

Similar to *D*_2_, in order to find out whether *D*_*st*_ has an inflection point (from increasing to decreasing), we make the following analysis. Consider two non-negative functions *f*_1_ and *f*_2_ are differentiable, the chain rule says,

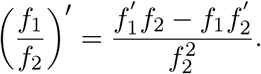

We set

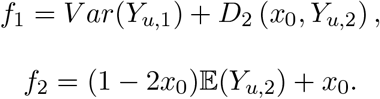

If 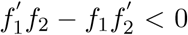, then (*f*_1_*/f*_2_)^′^ *<* 0. We extract the sign of (*f*_1_*/f*_2_)^′^ by 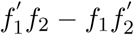. In the first step, the chain rule is used to find the approximate range of each parameter when the inflection point exists. In the second step, the parameters are selected according to the range to determine the approximate position of the inflection point. In the first step we consider the following limiting form as a case and give a result under *α <* 1 (see **Appendix** B.1),

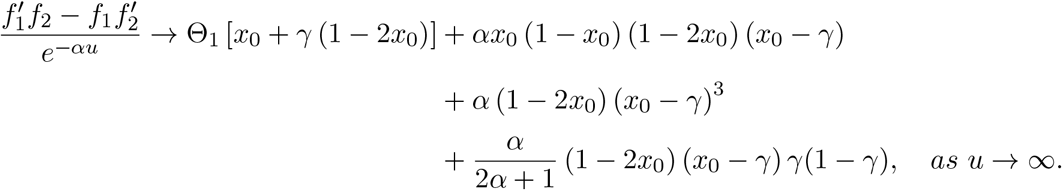

The above limit result contains *α* (Θ_1_ also contains *α*) and Θ_1_ contains (*x*_0_ − *γ*), which means that (1 − 2*x*_0_)(*x*_0_ − *γ*) determines the sign of all terms except Θ_1_*x*_0_ in the limit result. Consider fixing a positive value of *α*, then take the parameters *x*_0_ and *γ*, s.t. (1 − 2*x*_0_)(*x*_0_ − *γ*) *<* 0, and check the case where the limit is negative. Based on this process, we get the parameters *α* = 0.1, *γ* = 0.31 and *x*_0_ = 0.3 as a case where the inflection point should be (see Figure 5(a)). The above parameters were used to draw the curve of *D*_*st*_ with *u*. As shown in Figure 5(a), the inflection point does exist and is near 14.

**Figure 5:**
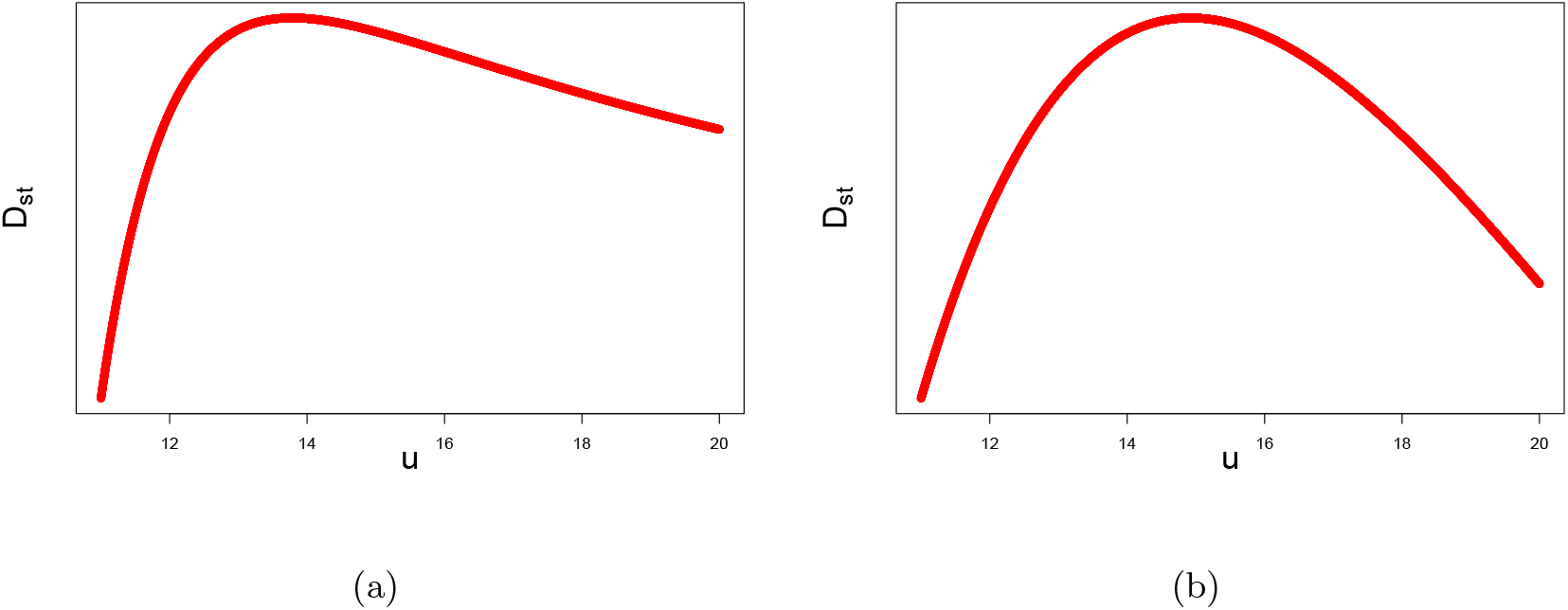
Shown is the curve of *D*_*st*_ with *u*. The inflection point does exist.

If we consider the same linear evolutionary pressure model for *P*_1_ and *P*_2_, then (18) can be simplified as

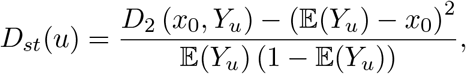

where,

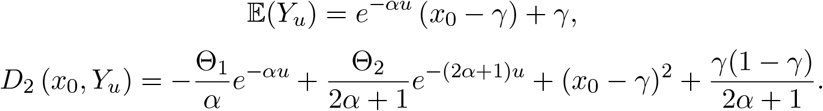

We set

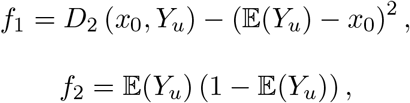

and take the limit to extract the sign part (see **Appendix** B.2),

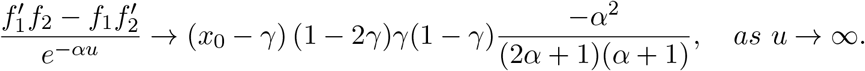

Obviously, we only need to consider *x*_0_ and *γ*, s.t. (*x*_0_ − *γ*) (1 − 2*γ*) *>* 0. We give the parameters, *α* = 0.1, *γ* = 0.4 and *x*_0_ = 0.6, to support our judgment (see Figure 5(b)). As shown in Figure 5(b), the inflection point is near 15.

## 3 Random migration rates

In the case of migration, we study the simplest linear form,

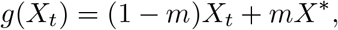

where the migration probability *m* is assumed to be fixed. In nature, however, populations migrate differently over time. Environmental climate, population size and other factors always affect the probability of migration in and out. Taking time dependence and randomness into account, we denote *m*_*t*_ as the migration probability, s.t.,

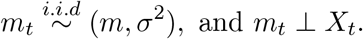

Note that the distribution of *m*_*t*_ is ignored and only its first two moments are marked as *m, σ*^2^ *>* 0. Assume 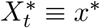, then

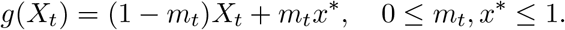

Following the Wright-Fisher model and approach taken in the previous section, we also give explicit formula for *F*_2_(*t*) and *D*_2_(*u*). The first two moments of *X*_*t*_ can be obtained,

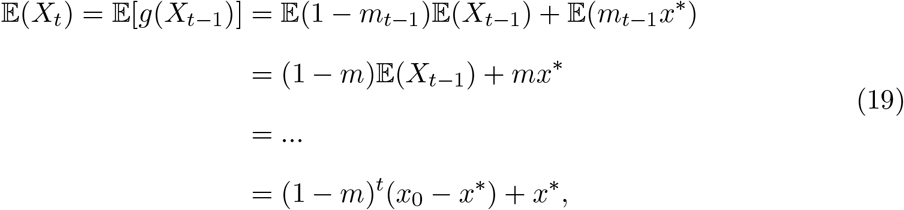

and

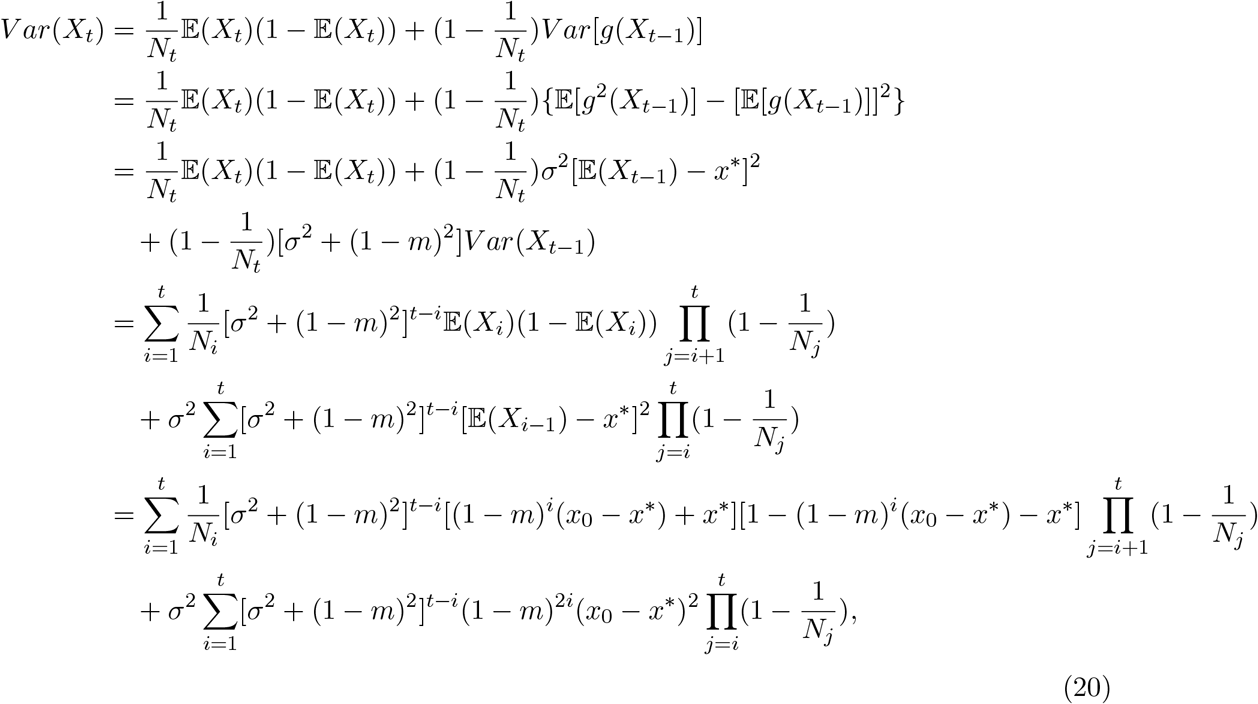

then by (19) we get

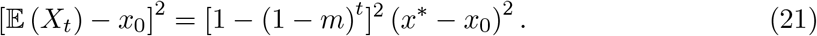

Combining (20) and (21), we follow the definition (6)

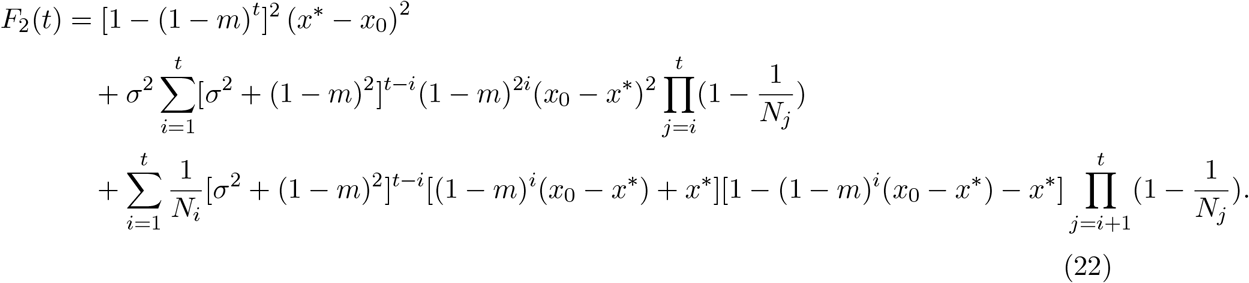

By scaling parameters *m* and *σ*^2^ in units of *N*_0_, we introduce

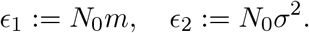

Following the previous definitions, we have

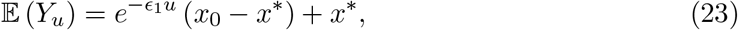

and

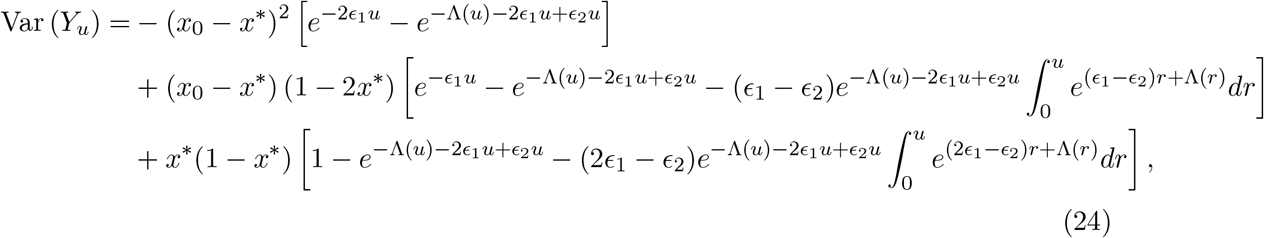

then by (23) we get

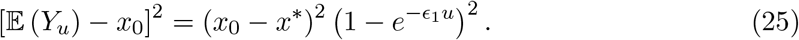

Combining (24) and (25), we follow the definition (10)

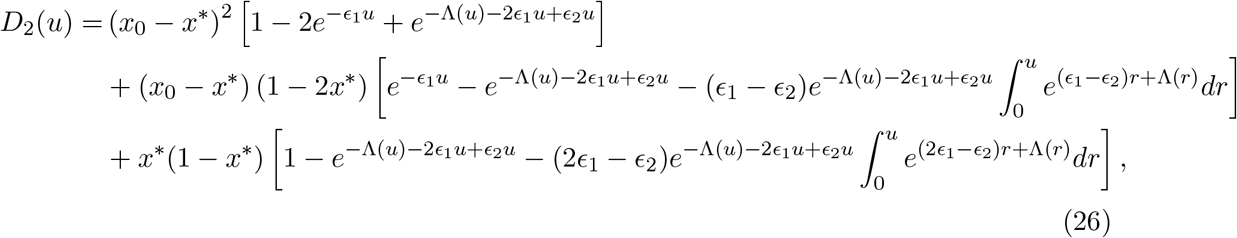

With the expressions of *F*_2_(*t*) and *D*_2_(*u*), we can use the method in the paper to reasonably discuss the parameters and the existence of inflection points. And the study of *F*_*st*_ with respect to *F*_2_, further research results will be carried out in our future work.

## 4 Discussion

We found expressions for the *F*_2_-statistic and the *F*_*st*_-statistic and how they vary over time. In general, we find that even for small population sizes, the behavior of the two statistics are well approximated by large population scaled expressions, considering time and paprameters scaled in units of population size.

Of particular interest is that migration might give rise to non-monotonic behavior. As real world most populations are subjected to migration, then this points to the conclusion that the behavior of the *F* -statistics for real world population in most cases will be non-monotonic.

It is worth mentioning that our proposed method only considers the first two moments of the allele frequency *X*_*t*_, which is also applicable to other cases as long as the second moment exists. In such a case, we can still give reasonable results for the linear representation of evolutionary forces, the expression of *F* -statistics and related parameters analysis. In population genetic studies, nonlinear factors such as natural selection are usually considered in order to explore the process of allele frequency change caused by evolutionary forces. In such a complex study, it is not hard to imagine that there would be no explicit expression for *F* -statistics. Therefore, the diffusion approximation of this setting is a mean of conducting similar analysis (Ewens, 2004; Lacerda and Seoighe, 2014; Tataru et al., 2015). We consider these as future research directions.

## Acknowledgements

The authors acknowledge the financial support from the funding agency of China Scholarship Council. The authors are supported by the Independent Research Fund Denmark (grant number: 8021-00360B) and the University of Copenhagen through the Data+ initiative.

## Appendix A Proofs

In appendix, we prove the theorems and corollary stated in the main text.

Random mating leads to a count of *A* alleles in generation *t* + 1 that is binomially distributed,

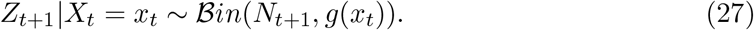

The goal is to account for effects of evolutionary and demographic forces on allele frequencies over time. We first calculate the first two moments of the allele frequencies.

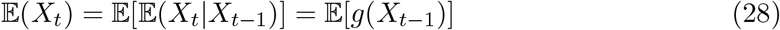

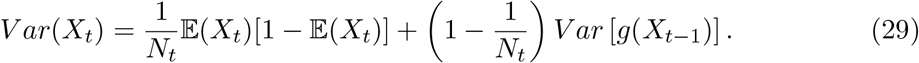

In the following, we treat the general linear case

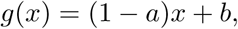

the mean and variance expression (28, 29) may be replaced by

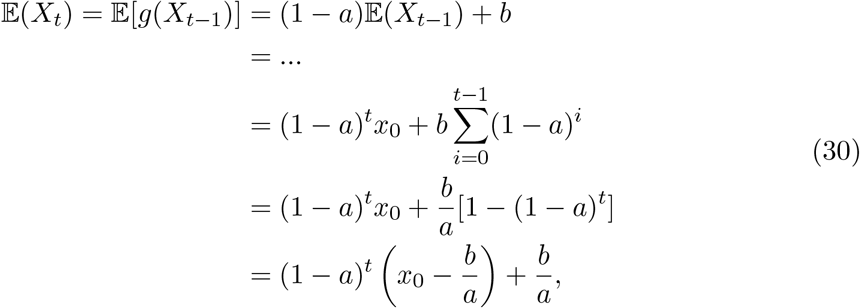

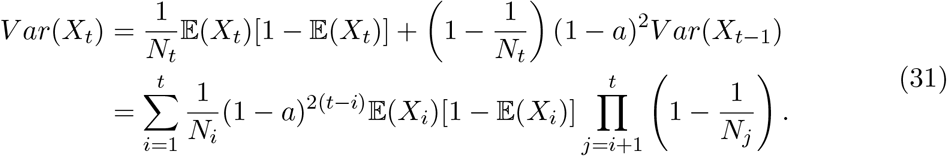

To consider approximations resulting from the infinite population limit, we take some appropriate variable transformations,

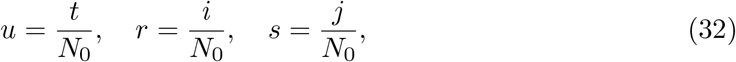

and

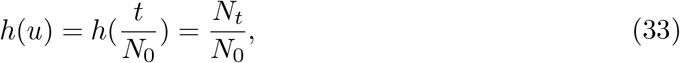

where *u, r, s* ∈ ℝ_+_, *i, j* = 1, …, *t* ∈ ℕ and *h* ∈ *L*^1^(ℝ_+_). Using the Riemann sum and Taylor approximation, let *N*_0_ be large enough, then

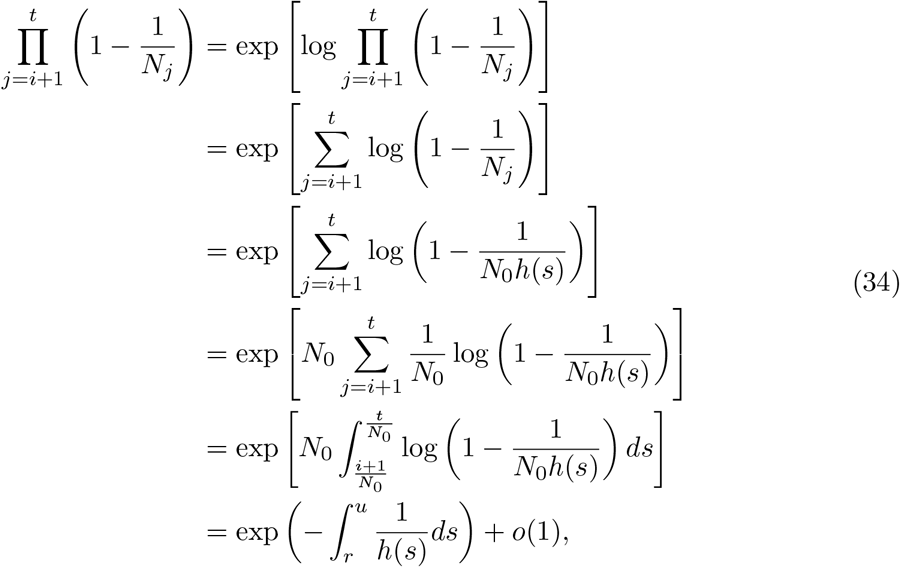

where *o*(1) is an infinitesimally small quantity, which is negligible in the limit. Using previous notations *α* = *N*_0_*a* and *β* = *N*_0_*b*, we have

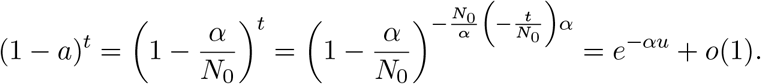

Defining 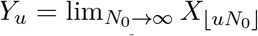 (Ewens, 2004), using (30),(31) and the Riemann sum, we obtain the mean and variance as a function of the scaled time

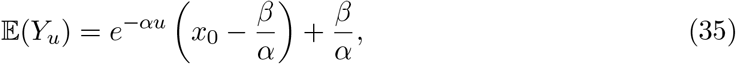

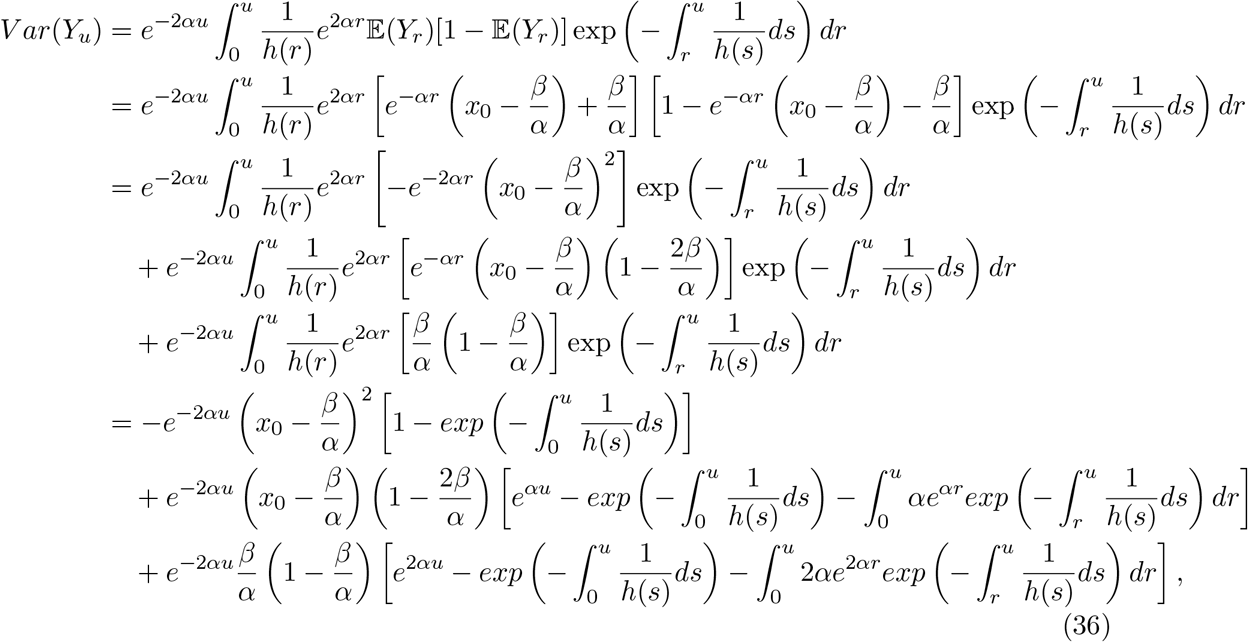

where, the last step in (36) was obtained using integration by parts.

### A.1 Proof of Theorem 1

*Proof*. Following expression (11), we consider the first derivative of *F*_2_ with respect to *t*,

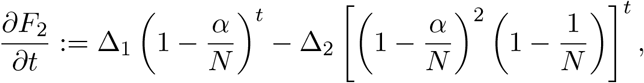

where

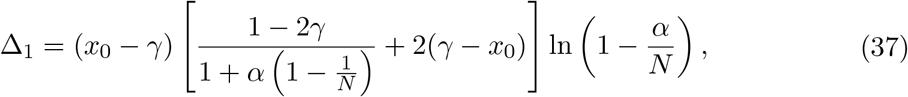

and

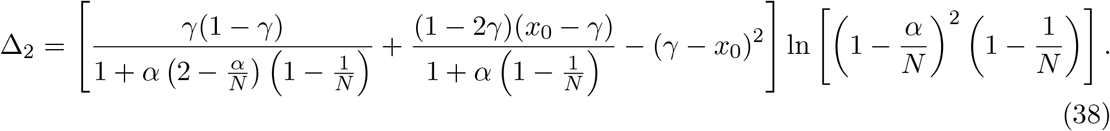

We want to observe where the inflection point of *F*_2_ occurs, so the following analysis is introduced,

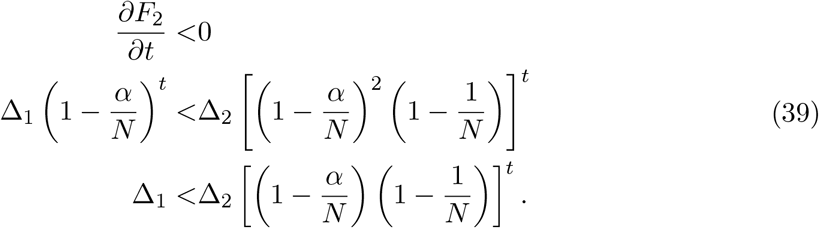

When Δ_2_ = 0, the last step in (39) indicates Δ_1_ *<* 0. However, by the definition (38),

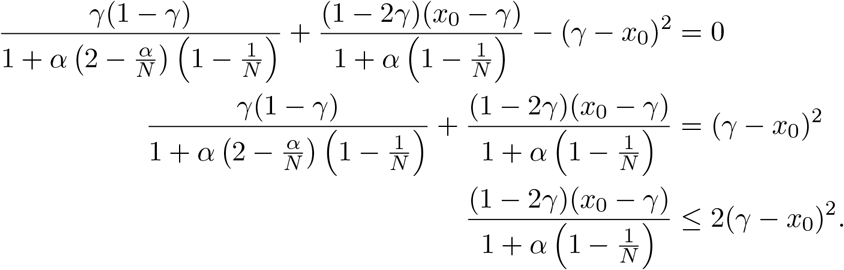

The last step was obtained using the following,

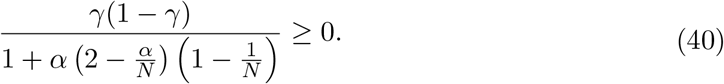

Hence, the definition (37) indicates Δ_1_ ≥ 0, that is a contradiction.

When Δ_2_ ≠ 0, (39) indicates

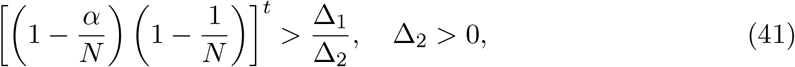

or

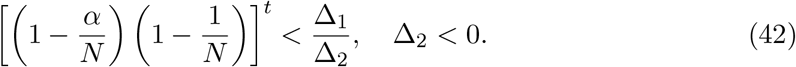

For (41), if Δ_1_ ≤ 0, by (40) and the definition (37),

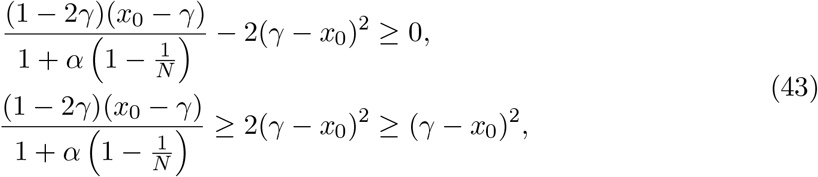

we get Δ_2_ ≤ 0, this contradicts Δ_2_ *>* 0. If Δ_1_ *>* 0 and 0 *<* Δ_2_ *<* Δ_1_, then 1 *<* Δ_1_*/*Δ_2_. However,

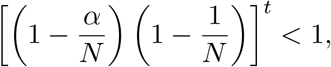

it contradicts (41). So, there may be an inflection point when 0 *<* Δ_1_ *<* Δ_2_, which makes *F*_2_ have a decreasing trend. In this case, using (41), we can get the range

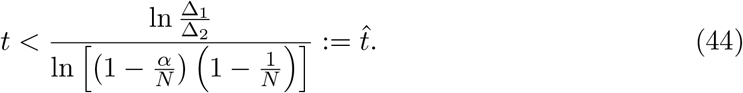

We know when *t* = 0 then *F*_2_ = 0 and *F*_2_ ≥ 0 for all *t*. 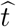 all depends on *N, α, γ, x*_0_ and 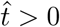 then 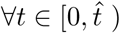, by (41) *∂F*_2_*/∂t <* 0, *F*_2_ (*X*_0_, *X*_*t*_) *< F*_2_ (*X*_0_, *X*_0_) = 0, that is a contradiction. From the above analysis, the existence of the inflection point of *F*_2_ means that (42) holds. For some *t >* 0, we prove

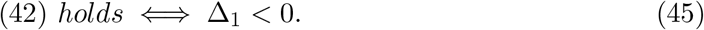

1. “=⇒” Consider proof by contradiction. Clearly, if Δ_1_ ≥ 0 then we get

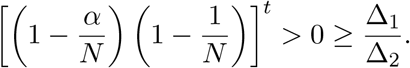
2. “⇐=” Clearly,

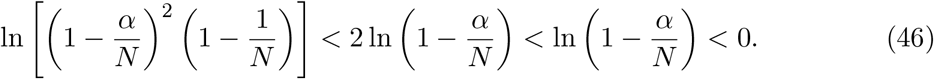

Using Δ_1_ *<* 0 and the definition (37), then (43) and (40) are satisfied. We obtain

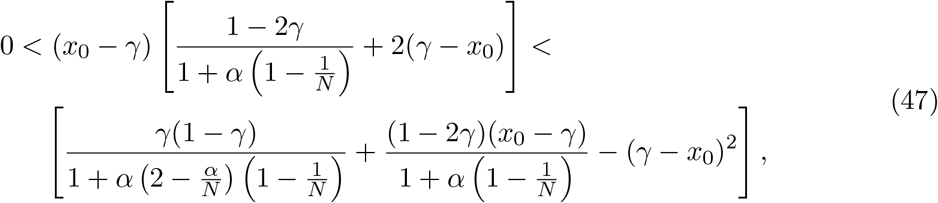

Combining (46) and (47),

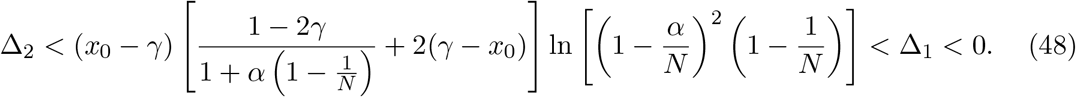

(48) indicates 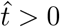 and for all 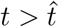, we have

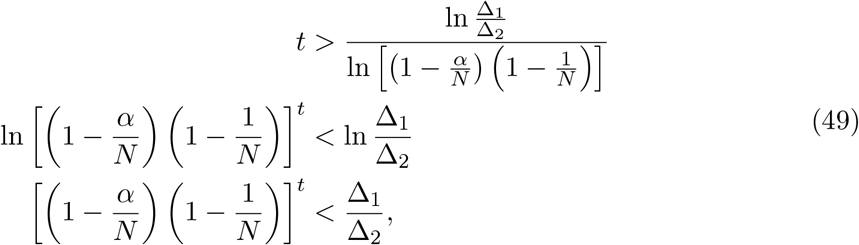

(42) holds.

All the above proof take into account that *F*_2_ is a continuous function of *t*, and even though we only use *t* ∈ ℕ, the conclusion still holds.

### A.2 Proof of Theorem 2

*Proof*. Following expression (12), we consider the first derivative of *D*_2_(*u*) with respect to *u*,

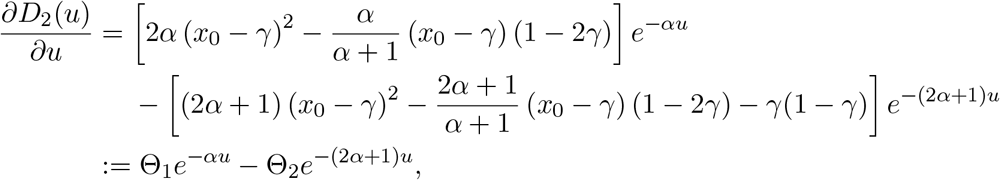

where,

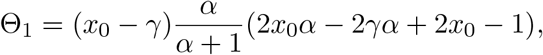

and

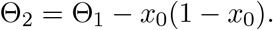

Refer to the proof of Theorem 1 (A.1), the following analysis is introduced,

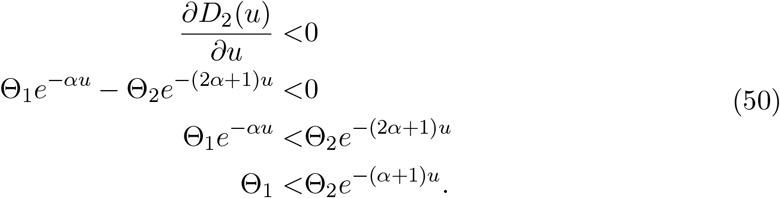

When Θ_2_ = 0, then Θ_1_ = *x*_0_(1 − *x*_0_) ≥ 0, that contradicts the last step indicating Θ_1_ *<* 0 in (50). When Θ_2_ ≠ 0, then (50) can be further transformed into

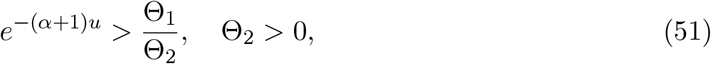

or

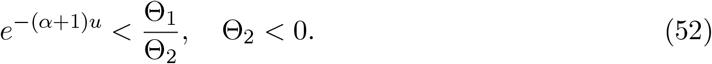

If Θ_2_ *>* 0 then Θ_1_ *>* Θ_2_ *>* 0, we have *e*^−(*α*+1)*u*^ *<* 1 *<* Θ_1_*/*Θ_2_. For (51), *D*_2_(*u*) cannot have an inflection point in its decreasing trend. If Θ_2_ *<* 0, Θ_1_ *>* 0, then Θ_1_*/*Θ_2_ *<* 0, that is not possible; if Θ_1_, Θ_2_ *<* 0 and obviously, Θ_2_ *<* Θ_1_ *<* 0 indicates Θ_2_*/*Θ_1_ *>* 1, that is possible. For (52), we can get the range,

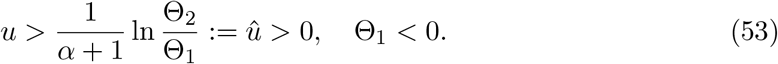

The process details are similar to the proof of Theorem 1 (A.1) and are partially omitted here.

### A.3 Proof of Corollary

*Proof*. According to Theorem 1 and 2, the necessary and sufficient condition for the existence of the inflection point is Δ_1_ *<* 0 and Θ_1_ *<* 0, respectively. To prove the corollary in 2.2.2, we introduce the following analysis.

For Δ_1_ or Θ_1_, we set *α, N, γ* as constants, then Δ_1_ or Θ_1_ is a parabolic function of *x*_0_. And for a parabolic function, the fact that existence of the roots depends on the sign of the discriminant, which is exactly that the discriminant Δ ≥ 0. In addition, since *x*_0_ ∈ [0, 1], we consider the range of the roots to complete the proof.

For Δ_1_, we know

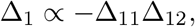

where, Δ_11_ = *x*_0_ − *γ* and

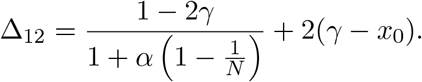

Then

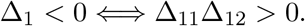

Defining Δ * = Δ_11_Δ_12_, obviously,

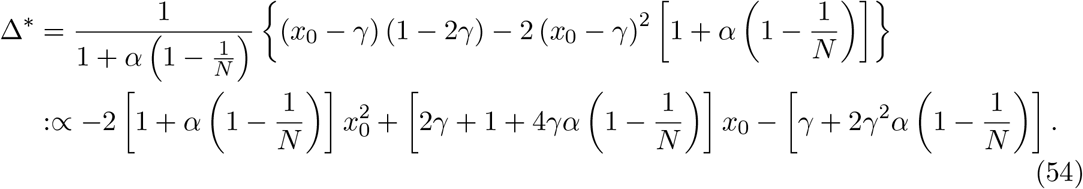

Using the last expression in (54), we calculate the discriminant

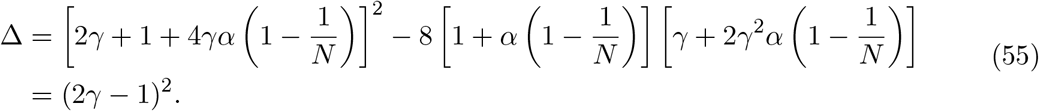

For ∀*α* ≥ 0, *N* ∈ ℕ, *γ* ∈ [0, 1] and *γ* ≠ 0.5, Δ *>* 0. And the expressions for the two roots are

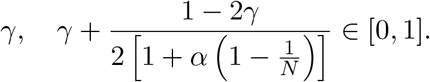

Hence, for a downward opening parabola, ∀*x*_0_ ∈ (*x*_0*L*_, *x*_0*R*_), where

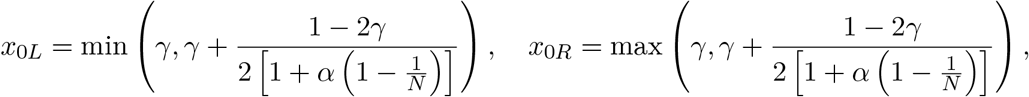

then Δ* *>* 0 and Δ_1_ *<* 0. According Theorem 1, *F*_2_(*t*) has an inflection point.

For Θ_1_, the process details are similar to the above steps and are omitted here.

## Appendix B The derivative and limit

For the *D*_*st*_(*u*) in 2.2.3, the inflection point is sought by the derivative and limit. We describe the calculations used in the formation of ideas as follows.

### B.1 Pure drift for *P*_1_, the linear pressure model for *P*_2_

We have

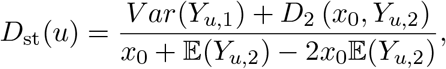

where

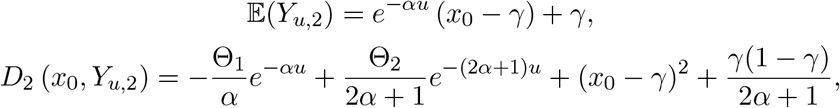

and

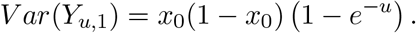

When two non-negative functions *f*_1_ and *f*_2_ are differentiable, the chain rule says,

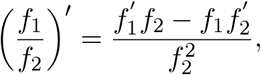

if *f*_1_^′^ *<* 0 and *f*_2_^′^ *>* 0, then (*f*_1_*/f*_2_)′ *<* 0. We set

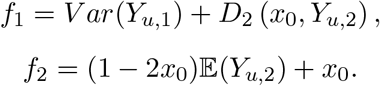

Using the above expressions, we get

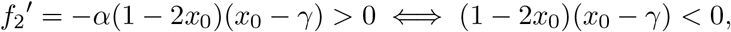

and

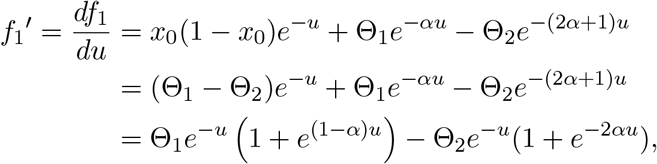

then

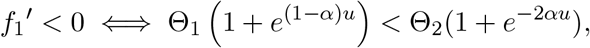

so,

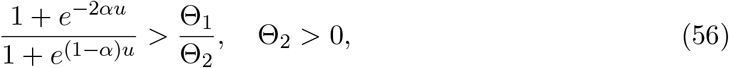

or

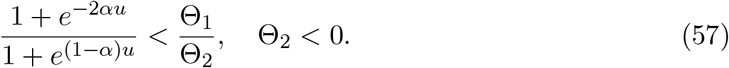

But from the previous analysis, we know that Θ_2_ *<* Θ_1_ and 0 *< e*^−2*αu*^ *< e*^(1−*α*)*u*^, (56) is not possible; (1 − 2*x*_0_)(*x*_0_ − *γ*) *<* 0 and Θ_1_ *<* 0 also cannot be held together in the areas delineated by the Figure 4, (57) is also undesirable. The above preliminary judgment inspires us to consider

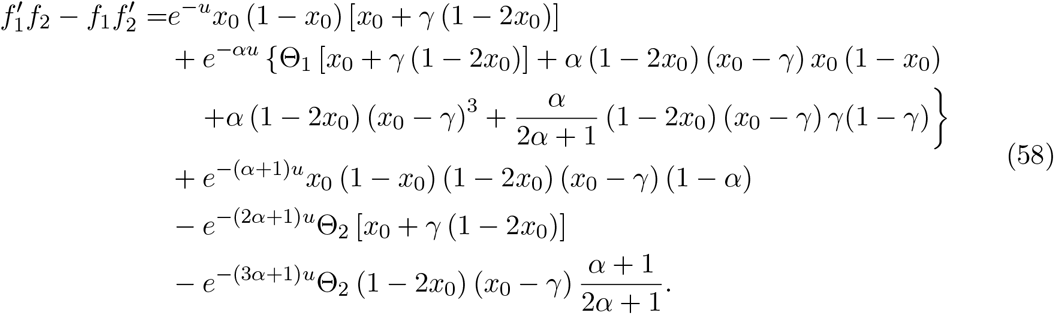

Letting *e*^−*u*^ = *λ*, then (58) can be transformed into

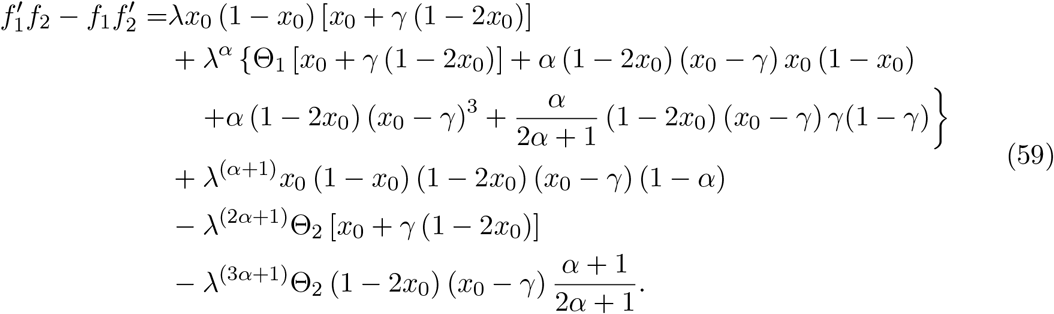

Using (59), we introduce the following limit.

1. If *α* > 1, *u* → ∞, then

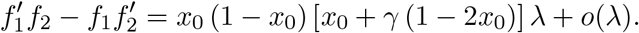
2. If *α* > 1, *u* → ∞, then

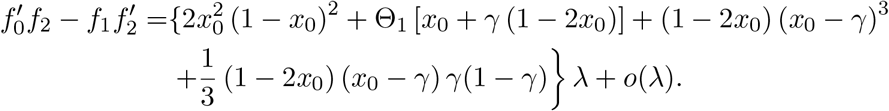
3. If *α* > 1, *u* → ∞, then

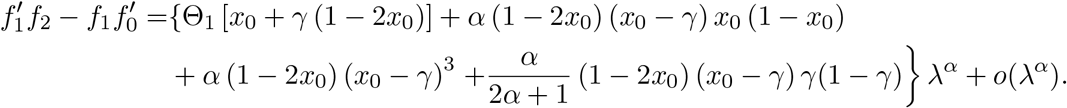

Obviously, in the **1)** 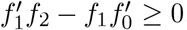, and 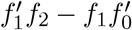 is more likely to be negative in the **3)** than in the **2)**. We consider the **3)** limiting form as a case and give a result under *α <* 1 in the main text.

### B.2 The same linear pressure model for *P*_1_ and *P*_2_

We consider

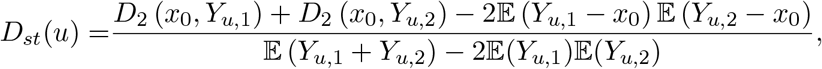

and the same linear evolutionary pressure model for *P*_1_ and *P*_2_, then *D*_*st*_(*u*) can be simplified as

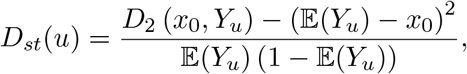

where,

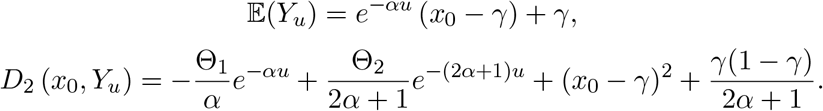

We set

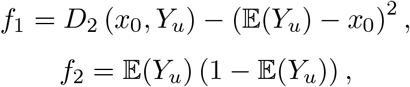

then in details,

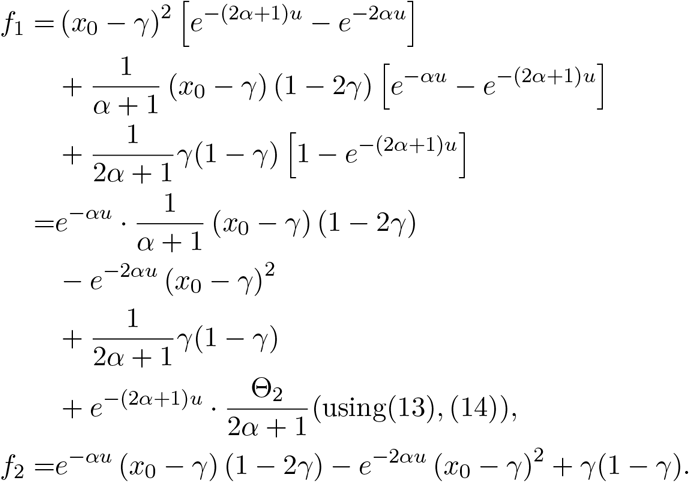

We calculate two derivatives,

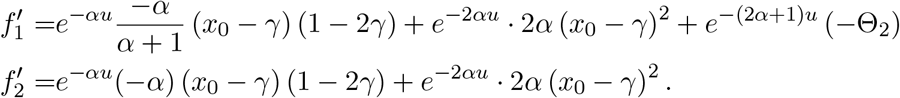

Combining 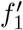 and 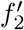, we obtain

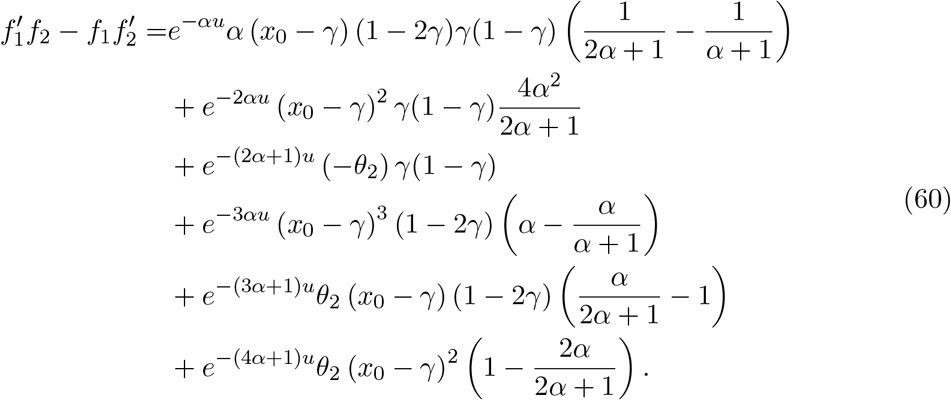

Letting *e*^−*u*^ = *λ* and *u* → ∞, then (60) can be transformed into

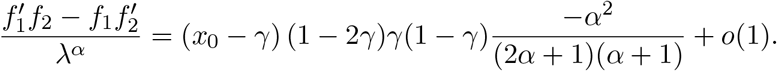

Obviously, we only need to consider *x*_0_ and *γ*, s.t. (*x*_0_ − *γ*) (1 − 2*γ*) *>* 0 and give a case in the main text.

